# Insulin-deficient diabetes impairs vaccine-mediated antibody and germinal center B-cell formation in mice

**DOI:** 10.1101/2025.09.24.677144

**Authors:** Christopher J. Genito, Pablo Ariel, Mark T. Heise, Lance R. Thurlow

**Author notes:** **Correspondence:** Christopher J. Genito, Lance R. Thurlow.

## Abstract

Individuals with diabetes are at increased risk for severe outcomes from vaccine-preventable infections and often mount weaker immune responses to vaccination. The factors underlying this impaired immunity remain unclear, and defining them is critical to improve vaccine strategies for this vulnerable population. Here, we focused on insulin deficiency as a contributing factor. Following immunization with an alum-adjuvanted protein subunit vaccine, insulin-deficient mice exhibited reduced antigen-specific IgG antibody responses, decreased lymphocyte numbers, and lower germinal center B-cell counts within the vaccine-draining lymph node. Three-dimensional whole-organ light sheet microscopy combined with virtual reality-assisted analysis revealed significantly smaller germinal center volumes in insulin-deficient mice than controls. These findings indicate that insulin deficiency can significantly constrain germinal center responses and impair antibody production from vaccination. Our results provide foundational evidence that diabetes-associated metabolic changes can significantly and negatively influence the quality of vaccine-induced immunity and highlight insulin deficiency as a potential physiological factor.

## 1. Introduction

Individuals with diabetes face a significantly higher risk for hospitalization and death from vaccine-preventable infections like influenza and COVID-19^1,2^. This elevated risk underscores the importance of vaccination to protect people with diabetes. However, the protection of this vulnerable population through immunization is complicated by reduced vaccine-mediated immune responses in both children and adults with diabetes, including responses from influenza^3^, COVID-19^4,5^, hepatitis B^6^, pneumococcal^7^, and MMR vaccines^8^. These deficits include lower production of protective antibodies^3,5–14^, reduced antibody longevity^4^, and impaired cellular responses^3,14–16^. Although the mechanisms remain unclear, poor glycemic control has been linked to weaker vaccine responses^3,16,17^, and uncontrolled diabetes is associated more broadly with immune dysfunction^18,19^. Type 1 and type 2 diabetes have been associated with decreased vaccine efficacy, suggesting a potentially shared mechanism. Both type 1 and type 2 diabetes are caused by impairments in insulin signaling, whether by insulin deficiency or insulin resistance. Insulin deficiency defines type 1 diabetes and can also arise in advanced stages of type 2 diabetes. Insulin therapy can restore some immune competence^20^, suggesting that insulin signaling itself may be an important determinant of the immune response to vaccines. We sought here to assess how insulin deficiency contributes to reduced vaccine responses in a well-established and experimentally tractable model of uncontrolled diabetes caused by the ablation of insulin-producing pancreatic beta cells.

Antibody responses elicited by immunization are major contributors to protection against most vaccine-preventable infections and are often used as key correlates of protection^21^. Following immunization, activation of naïve B-cells within B-cell follicles of vaccine-draining lymph nodes (LN) leads to the formation of germinal centers (GCs). Germinal center B-cells (GCBs) seed the B-cell response that gives rise to affinity-matured protective antibodies. A specific role for insulin in this process has yet to be fully elucidated, although insulin may influence both lymphocyte development and B-cell function. When insulin binding to its cellular receptor is inhibited, multipotent progenitors are skewed toward myeloid rather than lymphoid lineage differentiation^22^. Activated B cells also express the insulin receptor^23^. and downstream mediators of insulin signaling, including PI3K/Akt activation and glucose transporter 1 (GLUT1) expression, play important roles in B-cell activation^24^. In mice, insulin deficiency impairs antibody responses following vaccination^25–28^. However, the cellular mechanisms underlying this defect remain poorly understood. Because GCBs and GCs are the critical upstream drivers of affinity-matured antibody production, we hypothesized that insulin deficiency impairs vaccine-induced antibody responses by limiting GCB and GC responses.

A better understanding of the interplay between insulin deficiency and B-cell responses after immunization would be beneficial to inform vaccination strategies that may increase protection of people with diabetes from severe infections. Current strategies for subunit vaccines in clinical use and in preclinical development typically consist of a protein antigen in combination with an adjuvant to increase the strength of the immune response. For example, multiple COVID-19 vaccines are comprised of recombinant antigens based on the SARS-CoV-2 spike protein and the adjuvant alum^29^. Vaccine adjuvants, including alum, can enhance antibody responses to vaccines and facilitate the development of GCBs^30^. However, there is limited clinical evidence detailing how effective these vaccine strategies are specifically for people with diabetes, or how diabetes-related immune suppression or insulin deficiency affect immune responses to adjuvanted vaccines.

In the present study, we aimed to investigate how insulin deficiency influences vaccine-induced antibody responses and specifically examined GCB formation, an area where the effects of diabetes remain poorly defined. We therefore evaluated the ability of an alum-adjuvanted subunit vaccine to induce antibodies and GCB responses in insulin-deficient mice. Alum was the first FDA-approved adjuvant and currently remains one of the most used vaccine adjuvants. We used the well-characterized model protein antigen vaccine, ovalbumin (Ova), to test the impact of insulin deficiency on immune responses. We also used the streptozotocin (STZ) model of insulin-deficiency in mice, which specifically ablates the insulin-producing pancreatic β cells. Using the STZ model allowed us to interrogate the effects of insulin deficiency in the context of an immune system that is otherwise free of genetic or confounding metabolic defects. Mouse models also allow the enumeration and spatial analysis of GCBs within the vaccine-draining LN at a level of detail that is not feasible for study in humans. We used light sheet microscopy and virtual reality-based three-dimensional analysis to ensure the enumeration and volumetric determination of all GCs within the entirety of the lymph node. This modeled approach to vaccination in the context of diabetes led to several key findings of how insulin deficiency affects immune cell, GC structure, and antibody levels following vaccination. Our study demonstrates a clear connection between insulin deficiency and decreased antibody responses and GC formation from vaccination. These findings are foundational to an understanding of how diabetes affects the immune response to vaccination.

## 2. Materials and Methods

### 2.1. Animals and husbandry

Male and female C57BL/6J mice were acquired at 6-8 weeks old from the Jackson Laboratory and used in this study under an IACUC-approved protocol and housed in an AAALAC-accredited facility. Mice were housed in specific pathogen-free cages in groups of 5 with corncob bedding and environmental enrichment on a 12h/12h light/dark cycle and fed standard chow *ad libitum* with continuous access to water. Each individual experiment contained age-matched and sex-matched experimental groups, and the treatment/assessment order was arbitrarily assigned each day. At the study endpoint, mice were anesthetized with 500 mg/kg 2,2,2-tribromoethanol (Tokyo Chemical Industry) administered via intraperitoneal injection and euthanized by cervical dislocation while fully unconscious, as confirmed by loss of pedal withdrawal reflex. Experimental groups included vaccinated control mice, vaccinated insulin-deficient mice, naïve unvaccinated control mice, and naïve insulin-deficient mice.

### 2.2. Vaccine preparation and administration

Vaccines were formulated using ovalbumin (Ova) protein purified of endotoxin (Worthington Biochemical Corporation) and alum adjuvant in the form of 2% aluminum hydroxide gel (Alhydrogel, InvivoGen). Immediately before immunization, Ova was combined with alum in PBS to a final concentration of 40 µg/mL Ova and 1% alum. Mice were immunized once intramuscularly in the hind limb (gastrocnemius muscle) with 25 µL, corresponding to 1 µg Ova.

### 2.3. Generation of insulin-deficient mice

Mice were made insulin deficient by administration of streptozotocin (STZ). STZ was dissolved in sodium citrate buffer (0.1 M, pH 4) and injected peritoneally each day for 5 days consecutively at 90 mg/kg for female mice and 65 mg/kg for male mice. Mice were then given 7 days for diabetes onset before vaccination. Diabetes was confirmed if the blood glucose of each mouse reached ≥ 300 mg/dL before the day of immunization; otherwise, these mice were excluded from the study.

### 2.4. ELISA

Serum from mice was collected two weeks after immunization and analyzed for Ova-specific IgG antibody responses by ELISA. High-binding ELISA microplates (Grenier Bio-One) were coated with 500 ng/well Ova protein overnight at 4°C and then blocked for 1 h with dilution buffer (5% milk in PBS with 0.05% Tween20). Serum samples were diluted 20-fold in dilution buffer with additional 3-fold serial dilutions and added to wells for 2 h. HRP-conjugated goat anti-mouse IgG (H+L, human adsorbed, SouthernBiotech) or HRP-conjugated goat anti-mouse IgM (SouthernBiotech) was added at a 1:1000 dilution for 1 h. Plates were washed between each step three times with PBS + 0.05% Tween20, and then finally developed with 3,3’,5,5’-tetramethylbenzidine for 30 min. Developing was stopped with 50 uL 1M H_2_SO_4_. Optical density (OD) of developed plates was read at 450 nm with 570 nm background correction.

Titrations of OD vs. dilution were fit using GraphPad Prism 10 nonlinear fit (variable with four parameters). The background (bottom parameter) was set to the average value of negative control wells from the ELISA where no serum was added. Antibody titers were determined to be the interpolated dilution at which OD = 1. A positive control sample was included in each assay to account for day-to-day variation. Seroconversion was considered at any titer over the limit of detection (20, corresponding with the initial 20-fold sample dilution). Area under the curve (AUC) was also calculated for ELISA titration curves using Prism 10.

### 2.5. Spectral flow cytometry

Inguinal lymph nodes and spleens were removed from mice two weeks after immunization and processed for analysis by spectral flow cytometry. Tissues were collected into pre-weighed tubes with RPMI media supplemented with L-glutamine, 25 mM HEPES, 10% FBS, and 1% penicillin/streptomycin on ice. Tissues were then pressed through a 70 µm nylon filter. Spleen samples were treated for 2 min with ACK red blood cell lysis buffer at room temperature. Cells were then pelleted at 500 xg for 5 min, resuspended in media, and passed through an additional 40 µm nylon filter. A subset of isolated cells were stained with trypan blue and live cells were counted with a hemocytometer.

Cells were stained on ice with viability dye (50 ng/mL Pacific Blue-conjugated succinimidyl ester, Life Technologies) in PBS for 20 min, washed with flow cytometry staining buffer (Invitrogen), and blocked with TruStain FcX (BioLegend). Fluorescent antibody stain was performed for 30 min (antibodies are listed in **Table S1**) and cells were then fixed with BD Cytofix for 30 min. Cells were analyzed using a Cytek Aurora spectral flow cytometer with spectral unmixing performed using SpectroFlo software (Cytek). Gating and population analysis was performed using FCS Express 7 (De Novo Software).

### 2.6. Cytotoxicity assay

Direct toxicity of STZ on immune cells was determined by release of lactate dehydrogenase (LDH), measured using the CyQUANT LDH Cytotoxicity Assay Kit (Invitrogen). Immune cells were isolated from the spleens of naïve control mice as above and plated in a 96-well tissue cultured-treated round-bottom plate at a density of 2.5×10^5^ cells / 100 µL / well. Cells were incubated with STZ concentrations for 24 h at 37°C, 5% CO2. Viability was determined compared to untreated cells (representing 100% viability) and untreated cells lysed with the supplied lysis buffer at the 24 h endpoint (representing 0% viability).

### 2.7. Light sheet microscopy

Draining lymph nodes were fixed in 4% paraformaldehyde in PBS overnight at 4°C, then prepped, stained, and cleared for three-dimensional imaging using the Adipo-Clear method^31^, including embedding in 1% agarose. Antibodies used for staining are listed in **Table S1**. Samples were imaged using a light sheet microscope (LaVision BioTech UltraMicroscope II, Miltenyi Biotec) with immersion in dibenzyl ether. Emission filters (for lasers) were Chroma ET525/50m (488 nm), ET600/50m (561 nm), and ET690/50m (647 nm). Images were acquired using an Olympus MVPLAPO 2X/0.5 objective (with corrected dipping cap, 6 mm working distance) coupled to a zoom body and an Andor Zyla 5.5 sCMOS camera. Imaging parameter details are listed in **Table S2**. The 488 nm laser channel was used to capture autofluorescence.

### 2.8. Three-dimensional image analysis in virtual reality

Lymph nodes were virtually dissected from the surrounding adipose tissue by manually drawing a boundary at the 2D interface in XY planes and compiling a 3D surface using Imaris 10.2 (Oxford Instruments). The dissected LNs were then imported into syGlass v2.1 immersive virtual reality software operated using Meta Quest 3 mixed reality headset. Germinal centers were identified as GL7^+^ cells within follicles of B220^+^ or IgD^+^ cells using the “average” projection. GC borders were established by manually setting a threshold which clearly separated the GC from background fluorescence in the GL7 channel (detailed settings in **Table S3**). GCs were masked in syGlass and masks were then imported into Imaris for volume determination. Volume was determined using the continuous border around identified GCs.

### 2.9. Statistical analysis

Statistical analyses were performed using GraphPad Prism 10. Antibody titers, cell counts, and GC volumes were log-transformed before statistical analysis. Comparisons between three or more groups were corrected for multiple comparisons as denoted in figure legends. Preliminary power analysis determined group sizes of n=10 had power of >0.8 to determine a 2.5-fold difference in antibody titer and n=4 had >0.8 power to determine 2-fold differences in germinal center volume by light sheet microscopy.

## 3. Results

### 3.1. Minimal antibodies are elicited after vaccination in insulin-deficient mice

Insulin deficiency was modeled in mice to determine its effect on immunological outcomes from vaccination. Mice were made deficient in insulin production by administration of STZ, showing signs of diabetes onset by the end of the STZ treatment period and robust induction of diabetes via blood glucose levels one week later (**Fig. S1**). Diabetic insulin-deficient mice were then immunized intramuscularly with model vaccine antigen Ova and clinically relevant vaccine adjuvant alum (**Fig. 1A**). Two weeks after immunization, Ova-specific serum antibodies were measured by ELISA. Both titer (**Fig. 1B**) and area under the curve analysis of ELISA readouts (AUC, **Fig. S2AB**) showed that control animals reached significantly higher antibody responses than both insulin-deficient and vaccine-naïve animals. Insulin-deficient animals did not achieve anti-Ova IgG titers or ELISA AUC values that were significantly higher than naïve animals. In fact, only 2 of 13 (15%) of insulin-deficient mice displayed antibody seroconversion to the vaccine, defined by an IgG titer above the limit of detection (**Fig. 1C**). In contrast, 14 of 15 (93%) of control mice displayed seroconversion. We did not see strong signals of Ova-specific IgM by ELISA at this time point in either insulin-deficient or control animals (**Fig. S2CD**). We concluded that insulin-deficient mice were unable to form significant levels of antigen-specific IgG in response to an alum-adjuvanted vaccine.

**Figure 1.**
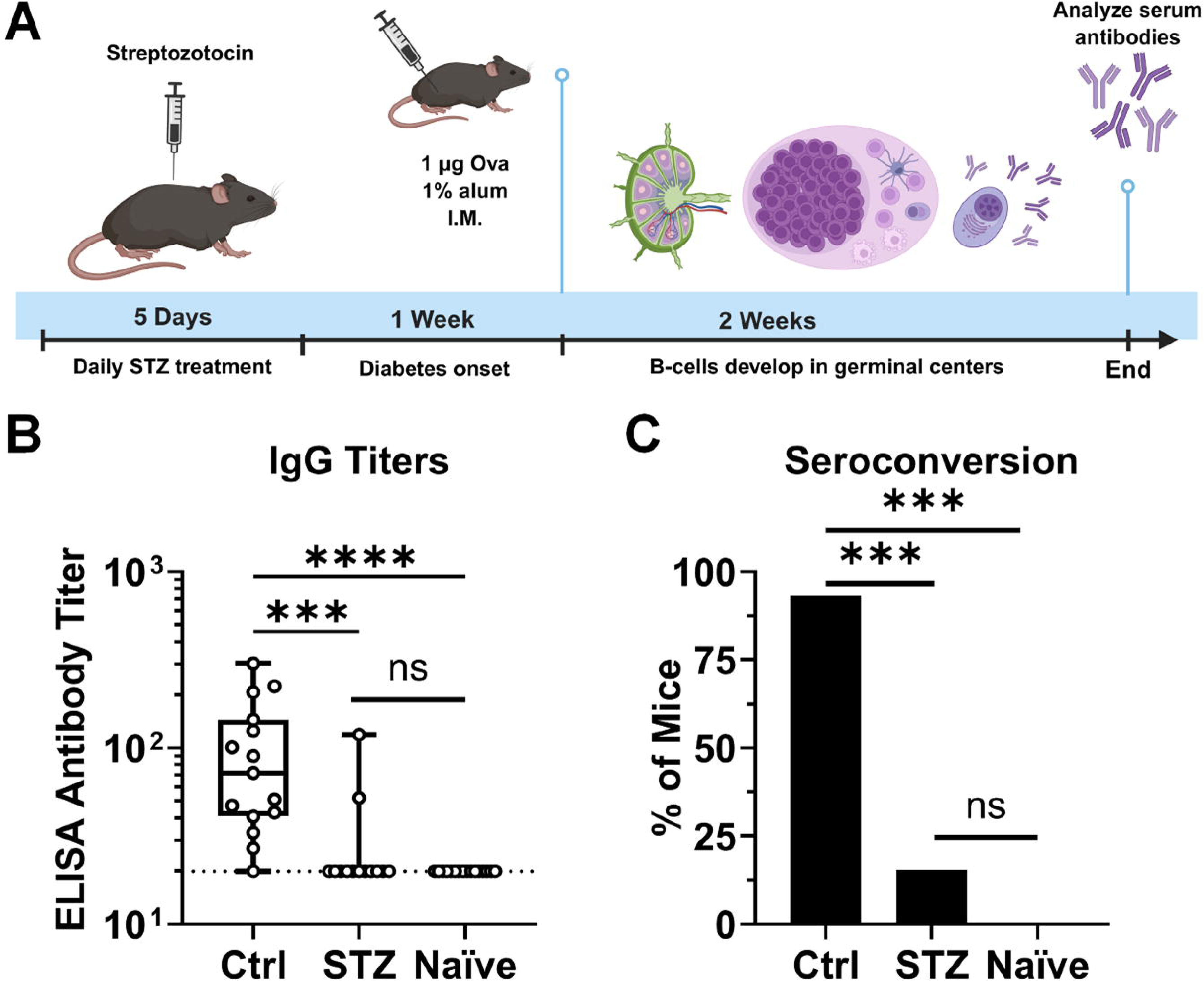
Antigen-specific IgG antibody levels are nearly absent in insulin-deficient mice after immunization with alum-adjuvanted vaccine. **(A)** Mice were made insulin-deficient using streptozotocin (STZ) and immunized with ovalbumin (Ova) adjuvanted with alum. **(B)** Two weeks after immunization, Ova-specific IgG antibody titers were quantified by ELISA for control (Ctrl; n = 15), STZ-treated (n = 13), and naïve (n = 15) mice. Data is presented as quartiles with a line at the median. Statistical comparisons were made using nonparametric Kruskal-Wallis test with Dunn’s correction for multiple comparisons. **(C)** Percentage of mice in each group that displayed seroconversion to the vaccine, defined as an IgG ELISA titer above the limit of detection. Statistical comparisons were made using Fisher’s exact test with Bonferroni’s correction for multiple comparisons. \*\*\**p* < 0.001, \*\*\*\**p* < 0.0001, ns = not significant. Data is pooled from three independent experiments.

### 3.2. Low lymphocyte numbers are observed within lymphoid tissue after vaccination in insulin-deficient mice

Vaccination with alum adjuvant causes swelling associated with responding immune cells proliferating in the draining lymph node (LN)^32^. We observed >2-fold smaller LN diameter from insulin-deficient mice vaccinated with Ova+alum compared to control animals (*p* < 0.0001, **Fig. 2AB**). In control animals, there was ∼3-fold increase (*p* = 0.001) in the number of live cells in the LN after vaccination compared to naïve animals (**Fig. 2C**). In contrast, this number was slightly decreased in vaccinated insulin-deficient animals compared to their naïve counterparts. We did not observe significant increases in weight or cellularity in spleens after vaccination, but there was a trend for fewer live cells and decreased weight for spleens from vaccinated insulin-deficient animals compared to vaccinated controls (**Fig. S3**).

**Figure 2.**
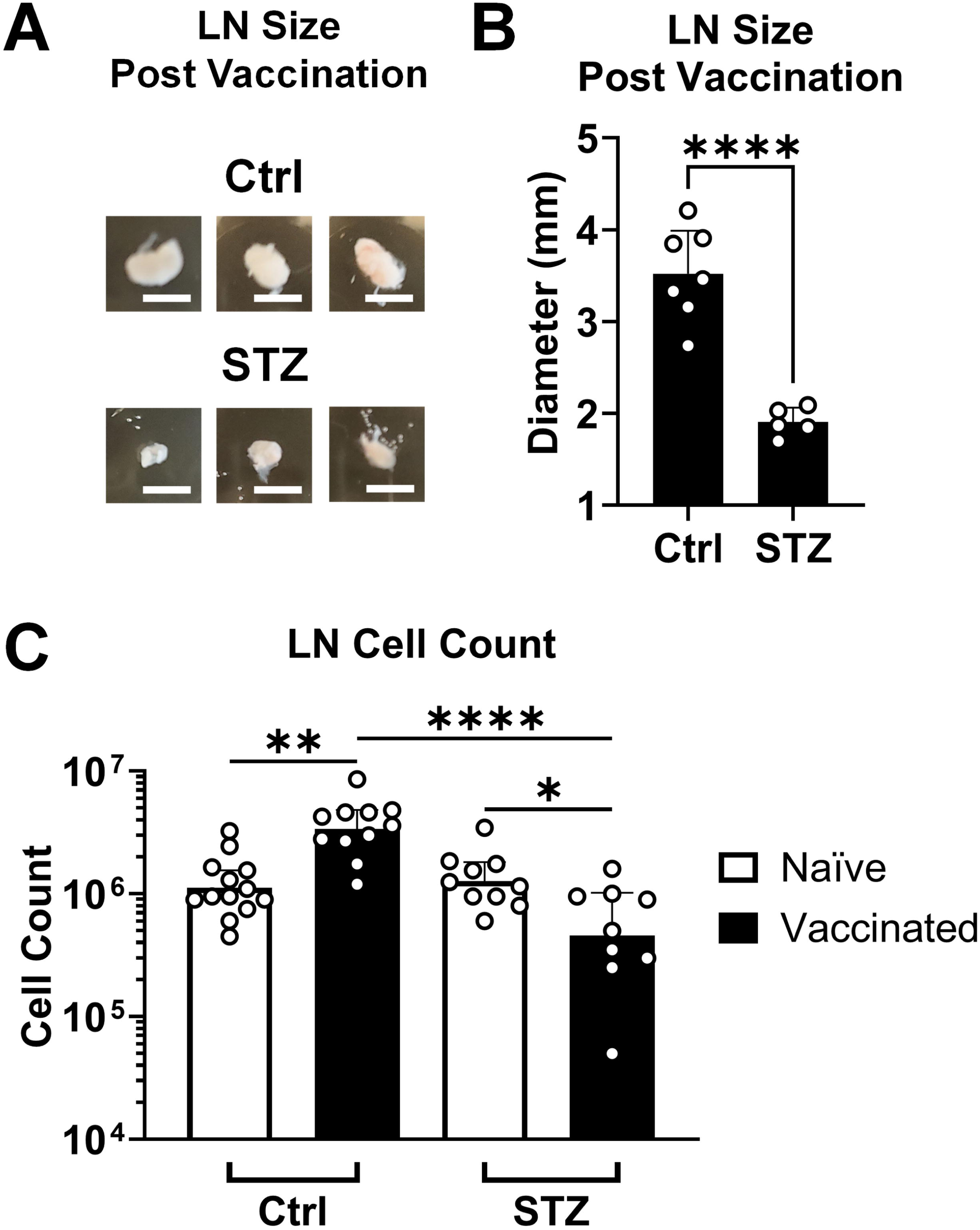
Draining lymph node size and cellularity after vaccination is significantly diminished in insulin-deficient mice. Draining inguinal lymph nodes (LN) were removed 2 weeks after vaccination with ovalbumin adjuvanted with alum from mice made insulin-deficient through streptozotocin administration (STZ) or from control animals (Ctrl). **(A)** Representative images of removed LNs. Scale bar = 3 mm. **(B)** LN size is quantified by measuring the semi-major elliptical axis (largest diameter) of each LN. Data are represented by mean + 95% CI and statistically compared by Student’s *t*-test (Ctrl n = 7, STZ n = 5). **(C)** Cells were isolated from LNs and live cell counts were determined by staining with trypan blue for Ctrl (naïve n = 13, vaccinated n = 11) and STZ-treated (naïve n = 10, vaccinated n = 9) mice. Data are represented by geometric mean + 95% CI and statistically compared by ANOVA with Tukey’s correction for multiple comparisons. \**p* < 0.05, \*\**p* < 0.01, \*\*\*\**p* < 0.0001 for all relevant comparisons. Data was pooled from 3 independent experiments.

The number of immune cells quantified in the vaccine-draining LN were consistent with the overall cell counts and LN sizes (**Fig. 2**): increased in control animals and slightly decreased in insulin-deficient animals after vaccination compared to their respective naïve counterparts (**Fig. 3A**). Both B-cells and T-cells, the largest immune cell general subpopulations, were proportional in number to the total immune cell count in draining LNs (**Fig. 3BC, Fig. S4**). There were no differences in B-cell or T-cell frequencies among total immune cells in the LN (**Fig. 3DE**). The fact that vaccinated insulin-deficient mice showed lower B- and T-cell counts than vaccinated control mice, but similar frequencies within the draining LN, suggested that neither immune cell population was disproportionately affected by insulin deficiency. Total immune cell counts in the spleen largely reflected the total number of live cells (**Fig. S3**, **Fig. S5A**). However, T-cells were more affected than B-cells by insulin deficiency in both number and frequency within the spleen (**Fig. S5BCDE**). Overall, it appeared that lymphocyte numbers in the relevant lymphoid organs were low after vaccination in insulin-deficient animals compared to naïve insulin-deficient counterparts, potentially contracting in vaccine-draining LNs. This was consistent with significantly higher frequencies of dead immune cell populations within the LNs of vaccinated insulin-deficient mice compared to vaccinated controls (**Fig. S6A**).

**Figure 3.**
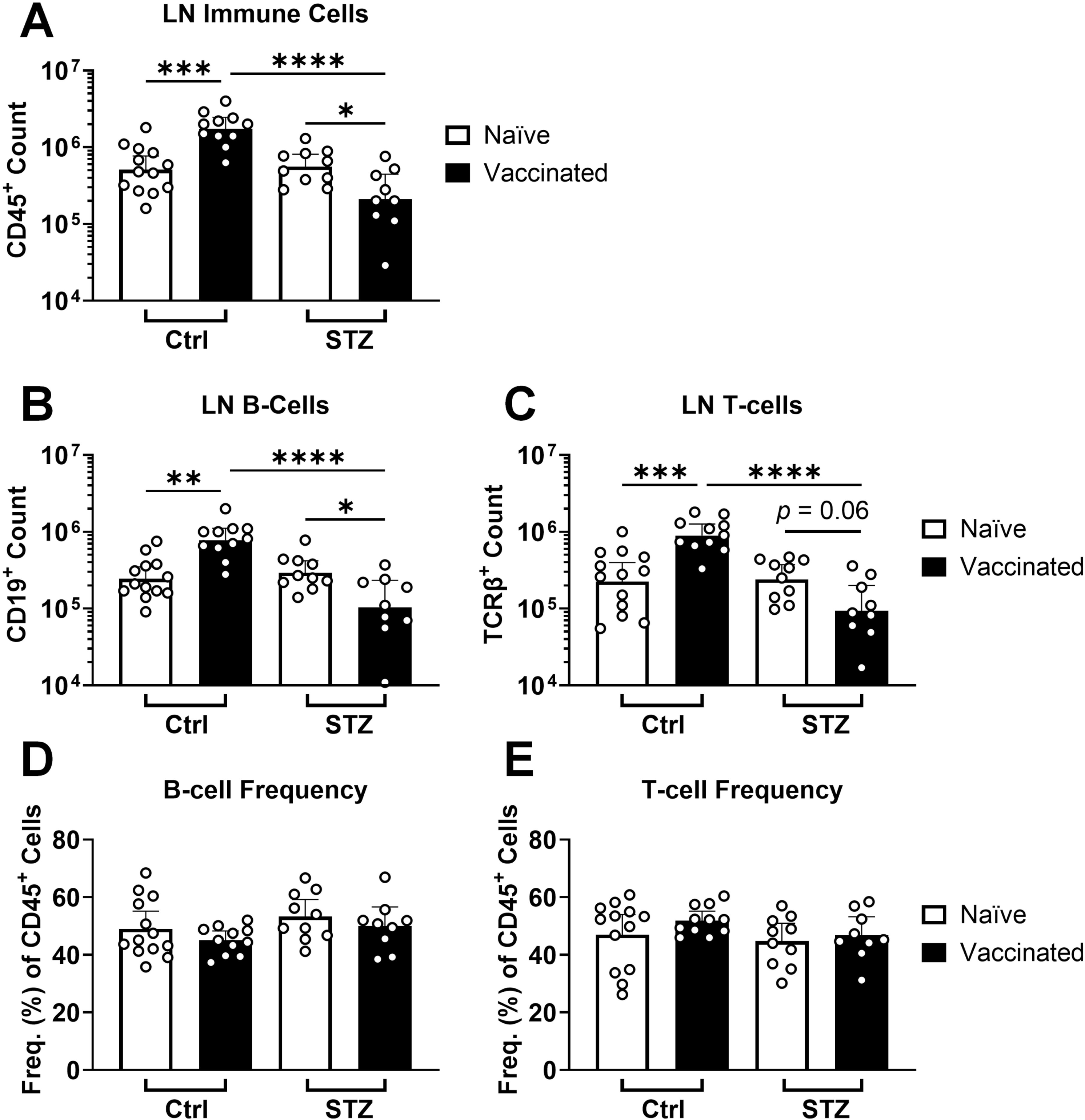
Vaccination fails to increase draining lymph node lymphocyte counts in insulin-deficient mice. Total immune cell **(A)**, total B-cell **(B)**, and total T-cell **(C)** counts, as well as B-cell **(D)** and T-cell **(E)** frequencies among total immune cells, were quantified by flow cytometry in vaccine-draining inguinal lymph nodes 2 weeks after vaccination with ovalbumin adjuvanted with alum for insulin-deficient (STZ, naïve n = 10, vaccinated n = 9) and control (Ctrl, naïve n = 13, vaccinated n = 11) mice. Data is represented as geometric mean + 95% CI. Statistical comparisons were made using ANOVA with Tukey’s correction for multiple comparisons. \**p* < 0.05, \*\**p* < 0.01, \*\*\**p* < 0.001, \*\*\*\**p* < 0.0001. Data was pooled from 3 independent experiments.

To test if observed decreases in immune cells of insulin-deficient mice were due to STZ-related toxicity, we isolated immune cells from the spleens of naïve control animals and incubated them with a range of doses extending well above published plasma concentrations reported for STZ administration in mice^33^ (**Fig. S6B**). Concentrations up to 2-fold higher than the highest reported STZ plasma concentration did not affect the viability of splenocytes after 24 h of co-incubation. Indeed, toxicity at >150-fold higher than the highest reported plasma concentration was only moderate (∼30%). Importantly, the maximum plasma concentration in mice was reported for a 200 mg/kg injection of STZ, which was 2-3-fold higher than the injection doses used in this study. There were also no differences in the frequency of membrane-permeable dead cells detected by flow cytometry in the LNs of naïve STZ-treated animals compared to naïve controls (**Fig. S6A**), revealing no obvious signal for STZ-related cell death within LNs. Overall, we concluded that decreases in cell counts observed in insulin-deficient mice were unlikely to be due to direct STZ-related toxicity on immune cells.

### 3.3. The germinal center response to vaccination is severely impacted in insulin-deficient mice

The frequency and number of germinal center B-cells (GCBs) within the draining LN were significantly increased 2 weeks after vaccination in control animals (**Fig. 4**). In stark contrast, the number nor frequency of GCBs were increased after vaccination in insulin-deficient mice, remaining at the level of naïve animals. Similar trends were observed in the spleen (**Fig. S7**).

**Figure 4.**
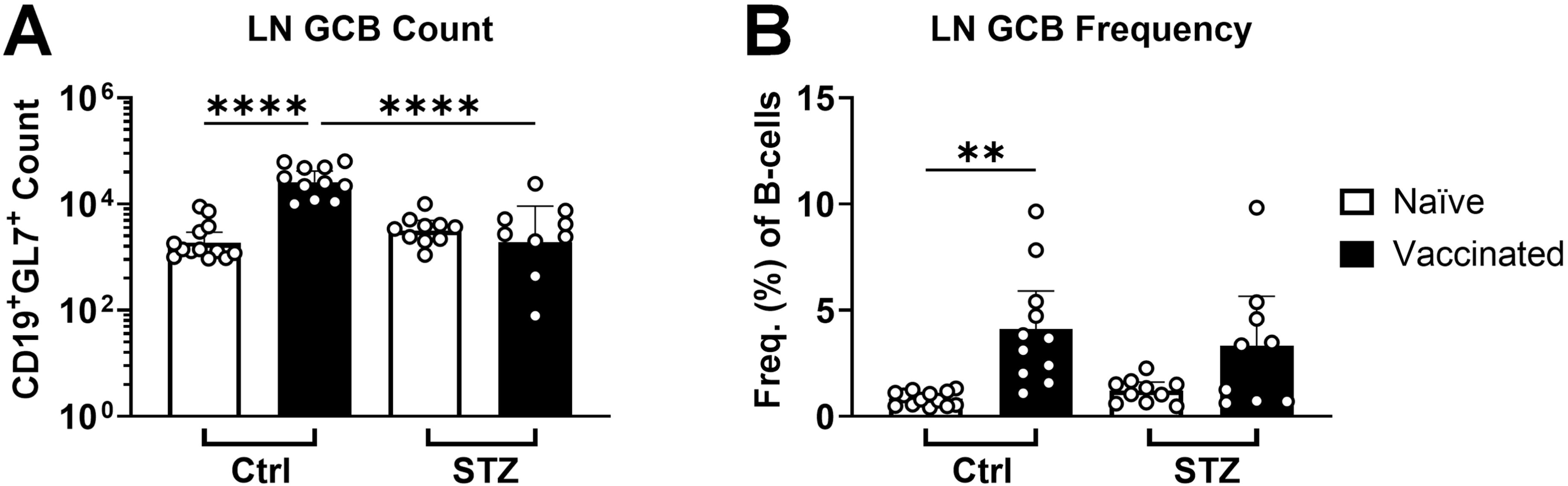
Vaccination fails to increase draining lymph node germinal center B-cell counts in insulin-deficient mice. GCB cell counts **(A)** and GCB frequency among B-cells **(B)**, were quantified by flow cytometry in vaccine-draining inguinal lymph nodes (LN) 2 weeks after vaccination with ovalbumin adjuvanted with alum for insulin-deficient (STZ, naïve n = 10, vaccinated n = 9) and control (Ctrl, naïve n = 13, vaccinated n = 11) mice. Data is represented as geometric mean + 95% CI. Statistical comparisons were made using ANOVA with Tukey’s correction for multiple comparisons. \*\**p* < 0.01, \*\*\*\**p* < 0.0001. Data was pooled from 3 independent experiments.

Given no detectable increase in GCB number, we hypothesized that germinal centers were not being formed within the draining LN of insulin-deficient mice following vaccination. We used light-sheet microscopy to identify and accurately count any GCs within the whole draining LN two weeks after vaccination (**Fig. 5**). GCs were able to be identified as GL7^+^ clusters of cells within B-cell follicles identified by IgD or B220 markers (**Fig. S8**). Light sheet microscopy allowed the construction of three-dimensional images within virtual reality-based software (syGlass). The three-dimensional borders of each GC were determined within the virtual reality environment for volume determination (**Fig. 6, Video 1, Video 2**). Surprisingly, we detected a similar number of GCs within the LNs of vaccinated insulin-deficient mice and vaccinated controls (**Fig. 6ABC**). However, GCs within insulin-deficient mice had significantly and drastically less volume (**Fig. 6DE**). The median GC volume was 10-fold less in the lymph nodes of insulin-deficient mice following vaccination compared to controls. We therefore concluded that the germinal center response to vaccination is severely impacted in insulin-deficient mice.

**Figure 5.**
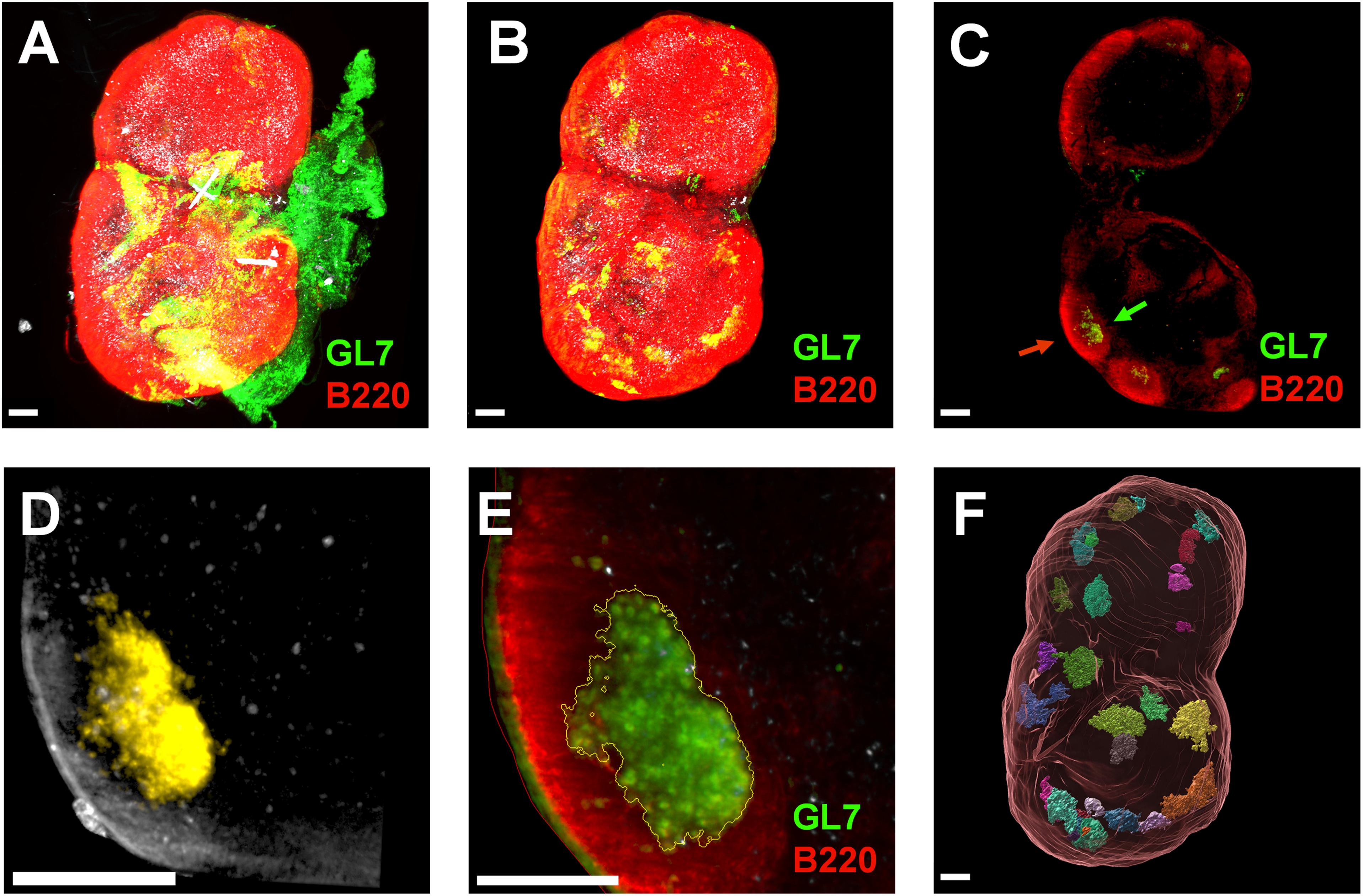
Virtual reality-assisted analysis enables confident detection and 3D modeling of germinal centers in whole lymph nodes imaged by light-sheet microscopy. **(A)** Three-dimensional rendering of draining inguinal lymph node (LN) 2 weeks after vaccination with ovalbumin adjuvanted with alum, imaged with light sheet microscopy (markers: red channel = B220, green channel = GL7; white channel = autofluorescence). **(B)** After manual identification of LN surface, the LN was virtually dissected from the surrounding adipose and connective tissue. **(C)** Representative two-dimensional slice of the 3D rendering in panel B. Germinal centers (GCs, green arrow) were identified as GL7^+^ clusters of cells within B220^+^ B-cell follicles (red arrow) within the LN. **(D)** Boundaries of identified GCs were established in virtual reality (syGlass) in the GL7 channel against a threshold background fluorescence. Cells included in the GC boundary are shown in yellow. **(E)** GC boundaries established in syGlass were imported into Imaris and processed as GC surfaces (yellow outline). The red outline denotes the LN surface. **(F)** Each opaque object (multicolor) is a 3D representation of a detected germinal center within the LN (translucent red outline). Scale bars represent 200 μm.

**Figure 6.**
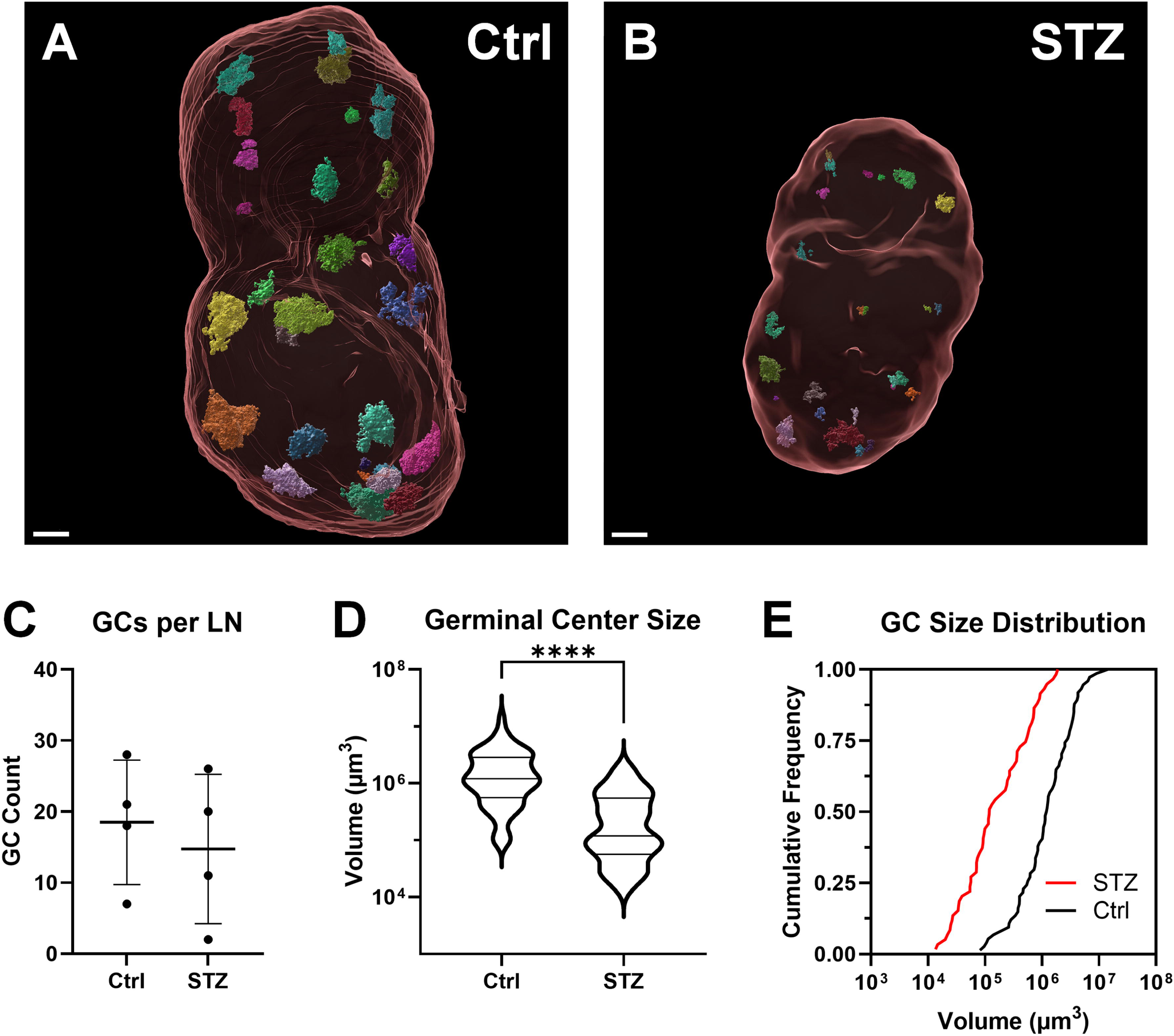
Germinal centers continue to form, but are 10-fold smaller, in insulin-deficient mice. Germinal centers (GCs) within draining inguinal lymph nodes (LN) were analyzed by light sheet microscopy 2 weeks after vaccination with ovalbumin and alum adjuvant in mice made insulin deficient by administration of streptozotocin (STZ). **(A,B)** Representative 3D images of LNs (red outline) and GCs (multicolor, opaque) are shown for control (Ctrl) and STZ-treated mice. **(C)** The total number of GCs in each lymph node (n = 4 LNs per group). Data are represented as mean±SD. **(D,E)** Cumulative size distribution of GC volumes (Ctrl, n = 74; STZ, n = 59). Data is represented as quartiles with line at median (D) and a frequency of distributions (E). \*\*\*\**p* < 0.0001, Kolmogorov-Smirnov non-parametric test for cumulative distributions. Data was pooled from 4 independent experiments. Scale bar = 200 μm.

## 4. Discussion

Poorly controlled diabetes increases the risk of severe outcomes from infection^1,2^ and has also been associated with poorer immune responses to vaccination to protect from these infections^3,4,6–17^. Our study expands on findings by others that have shown a deleterious effect of insulin deficiency on antibody formation from vaccination^25–28^. In addition to establishing a tractable model using a well-established model protein antigen and clinically relevant adjuvant, we have expanded the understanding of how insulin deficiency impairs humoral responses by demonstrating how insulin deficiency profoundly alters the GCB response to vaccination (**Fig. 4**) as well as structural GC responses using advanced whole-LN imaging (**Fig. 5**, **Fig. 6**). Quantifying GCBs by flow cytometry or immunofluorescence is straightforward, but these approaches provide limited information on GC architecture, especially at the whole LN scale. Conventional 2D histology introduces variability, as only a few sections from a larger 3D volume are analyzed. Confocal z-stacks, while enabling 3D reconstructions, can only do so for relatively small volumes, and are very labor-intensive for whole-LN analyses. By contrast, light-sheet microscopy permits rapid and unbiased volumetric imaging of all GCs within an LN. Although this approach has been applied in other contexts, its use in diabetes research has been limited. This is particularly relevant because diabetes-associated microvascular changes may alter LN architecture and GC organization, making whole-LN 3D analysis critical for detecting differences in GC number and volume that 2D sampling might miss. Moreover, because GC boundaries are highly dynamic and not sharply defined, immersive virtual reality-assisted segmentation can accelerate boundary detection and spatial mapping compared to conventional tools.

Using this approach, we found that vaccinated insulin-deficient mice generated similar numbers of GCs in vaccine-draining LNs compared to controls (**Fig. 6C**), but they were 10-times smaller (**Fig. 6DE**), in agreement with 10-fold lower GCB numbers (**Fig. 4A**). In insulin-deficient mice, GCB numbers did not increase with vaccination past the level of naïve controls (**Fig. 4A**), consistent with the idea that without insulin, a critical growth factor, the proliferative potential of GCBs and expansion of the GC may be impaired. The apparent contradiction between GC formation in insulin-deficient mice (**Fig. 6**), yet no increase in GCB abundance compared to naïve controls (**Fig. 4A**), may be explained by the observed small but significant contraction in total B-cell numbers upon vaccination (**Fig. 3B**). Regardless, GCs still formed in vaccinated insulin-deficient mice, but the observed 10-fold difference between insulin-deficient and control mice in GCB formation following vaccination (**Fig. 4A**) and 10-fold smaller GC volume (**Fig. 6**) strongly suggests an inability for GCB to expand in insulin-deficient mice. This contrast underscores the value of three-dimensional imaging approaches for resolving GC structure and volume and providing essential contextual information beyond cellular quantification. Although the precise mechanisms that contribute to B-cell contraction, lack of GCB abundance, and reduced GC volume are unknown, they could be a result of metabolic constraints, hyperglycemia, or related stressors. Furthermore, reduced LN size (**Fig. 2**, **Fig. 6**) may also impose spatial limitations on GC expansion.

Our data demonstrate that insulin deficiency constrains GC formation at both the cellular and structural level, revealing a previously unexplored mechanism by which impaired metabolic control may compromise vaccine responsiveness. For example, the observed impairment in GC formation likely impairs the formation of antibody-secreting plasma cells, which would explain the lower antibody responses observed after vaccination (**Fig. 1**). At two weeks post vaccination, we observed this impairment at a time point that represents peak GC formation. The formation of a similar number, yet smaller, GCs in the draining LNs of STZ-treated mice compared to controls (**Fig. 6**) provides the possibility that peak GC formation is delayed during insulin deficiency. However, delayed antibody responses from vaccination are not observed in people with diabetes, who instead experience faster rates of antibody decay^4^.

Our study importantly incorporates clinically relevant contexts by using a vaccine formulated with alum and administered through the intramuscular route, consistent with many licensed human vaccines. Despite this, insulin-deficient mice mounted weaker antibody and GCB responses (**Fig. 1**, **Fig. 4**, **Fig. 6**), indicating that standard vaccination strategies may be insufficient under conditions of metabolic dysregulation. Some clinical analyses of “simple” subunit vaccines without adjuvants show comparable immune responses elicited in people with or without diabetes^14,34,35^. However, more advanced vaccine platforms, including adjuvanted protein subunit and mRNA vaccines, have repeatedly revealed impaired immunogenicity in people with diabetes^4,5,14^. These discrepancies raise the critical question of whether platforms that enhance responses in the general population are equally effective in individuals with diabetes. Given that people with diabetes consistently experience worse outcomes following infection, developing vaccines that are specifically effective in this population is of pressing importance.

Using a model of STZ-induced diabetes in mice is advantageous to focus study on the effects of insulin deficiency on the immune system, allowing a controlled induction of diabetes in adult mice with otherwise intact immune systems. Other mouse models contain confounding immunological factors, such as autoimmune defects in non-obese diabetic (NOD) mice, perturbations of immune development resulting from early-onset diabetes in *Ins2^Akita^* mice, or obesity- and hormone-related effects (e.g., leptin) in other models of diabetes. While insulin deficiency specifically impairs immune function in the STZ model^36^, it has been historically suspected that STZ-related toxicity may also impact long-term splenic, thymic, and bone marrow immune cell viability in mice. In our model, mice were given a low dose of STZ to reduce the chance of toxicity outside of pancreatic β cells. Low dose regimens significantly reduce off-target toxicity^37,38^. The modest ∼6% weight loss we observed in animals during the STZ administration period corresponded with the onset of diabetic hyperglycemia and therefore cannot be confidently attributed to drug-related toxicity (**Fig. S1A**). Importantly, STZ was not directly toxic to splenic immune cells at physiological doses (**Fig. S6B**). We observed neither a reduction in B-cell or T-cell populations nor an increase in dead immune cells in the lymph nodes of naïve STZ-treated mice compared with naïve controls (**Fig. 3, Fig. S6A**), arguing against a substantial contribution of off-target STZ toxicity to the lymphocyte phenotypes observed in this study. Vaccines were given a week after the last STZ dose to ensure STZ presence did not influence immune cell activation by immunization. STZ is rapidly cleared by the kidneys and has a plasma half-life of less than 10 min in rodents^39^. In the absence of *in vivo* clearance mechanisms, STZ can persist *in vitro* for extended periods (up to 14 days in solution^40^); under these conditions, immune cell viability after 24 h exposure to concentrations higher than physiologically relevant (**Fig. S6B**) provides strong evidence against direct lymphocyte toxicity. Our study therefore did not contain evidence to directly link STZ toxicity to the observed deficiencies in the immune response to vaccination. Additionally, our observations of decreased B-cell and T-cell abundance after vaccination complement a previous report of reduced lymphocyte formation during viral infection in both STZ-treated mice and a non-pharmacologic model of insulin deficiency (*Ins2^Akita^* mice)^41^. Consistent with this prior evidence linking insulin deficiency to impaired lymphocyte responses, our finding of no detectable STZ-associated immune cell toxicity both *in vitro* and *in vivo* supports the interpretation that insulin deficiency is the primary driver of the observed phenotype in our model. Nonetheless, a partial effect of direct STZ action on immune cells or off-target effects that may indirectly affect immune cell function cannot be definitively ruled out.

Sufficient insulin signaling could be directly and/or indirectly required for B-cell activation. While insulin signaling has direct cellular involvement in T-cell expansion, metabolism, chemotaxis, and effector functions, the direct activity of insulin in B-cell activation and development remains unclear^24^. Here, we observed lower cell counts in both the T-cell and B-cell populations within the vaccine-draining LN after immunization in insulin-deficient mice compared to controls (**Fig. 3**). Cell-to-cell contact and cytokine production from T-cells, follicular dendritic cells, and other immune cells are important for GCB development within lymphoid tissues. In particular, T follicular helper cells (TFHs) are crucial mediators of GCB survival, activation, and development. Functional inhibition of TFHs could very well affect GCB responses. The formation of GCs in response to vaccination suggests that TFHs are engaged in our model, even in insulin-deficient mice (**Fig. 6**). However, whether TFHs rely on insulin signaling for their function to the same extent as conventional T-cells remains undescribed by previous studies, and whether insulin deficiency impairs TFH function was not established by these experiments. Another important consideration is that insulin may support immune function by preserving a favorable milieu for immune cell activation. Systemic physiological changes incurred by insulin deficiency potentially have significant effects on immune function in response to vaccination. For example, hyperglycemia can impair T-cell function including cellular trafficking, control of infection, and memory responses^42^, though the specific effect of hyperglycemia on B-cells is much less described. We observed hyperglycemia throughout the study period (**Fig. S1B**), making high glucose levels a potential contributing factor to decreased immune responses to vaccination. The interaction of insulin deficiency and the immune response to vaccines is likely multifaceted, and further study would be needed to elucidate if the impairments are due to a lack of insulin signaling on the immune cells themselves, or indirectly through intercellular interactions or maintaining a favorable metabolic environment for vaccine-mediated immunity. Because our model isolates insulin deficiency as the cause of diabetes, we therefore did not explore how other factors and co-morbidities related to diabetes like obesity, cardiovascular disease, and impaired wound healing may impact vaccine effectiveness. However, using this focused model demonstrated that insulin deficiency can potentially have a profound effect on GCB responses and antibody formation from vaccination.

The overall finding of this study is that the ability of an alum-adjuvanted protein subunit vaccine to elicit antibody and GC responses was markedly impaired in a mouse model of insulin deficiency. In particular, both GCB numbers and GC volume were reduced in vaccine-draining lymph nodes. A principal advance of this work is the development of a method to quantify changes in the 3D structure of GCs, which still form in insulin-deficient mice but at significantly reduced volume. Importantly, our work establishes a virtual reality-assisted whole-LN approach for identifying and analyzing GCs in three dimensions, providing a powerful tool for investigating how diabetes and other diseases alter spatial immunological processes within lymphoid tissues. We used a model subunit vaccine to interrogate these fundamental aspects of humoral immunity, which are of crucial importance considering that the majority of licensed vaccines confer protection primarily through the induction of antibody responses. Future work will address how these defects in humoral immunity translate to defects in vaccine-mediated protection from infection. This study is a step to understanding the role of insulin signaling in acquired immunity and gives context to future study on disease states where insulin signaling is impaired through resistance, most pertinent in type 2 diabetes and obesity. It nonetheless remains unclear how insulin resistance may specifically affect immune responses to vaccination and to what extent it might differ from insulin deficiency. However, these findings highlight insulin deficiency as a potential contributor to the defective vaccine responses observed in people with diabetes and underscore the need to define how diabetes-specific physiological factors shape vaccine-induced immunity. Elucidating the mechanisms underlying impaired GC formation and antibody production will be essential for optimizing vaccine design and tailoring immunization strategies for this vulnerable population. Such efforts may inform adjuvant selection, dosing strategies, or even the development of next-generation vaccines that account for the unique immunological considerations of people with diabetes.

## 6. Author Contributions

CJG, MTH, and LRT designed the studies. CJG and PA designed the data analysis methods. CJG carried out the experiments and analyzed the data. CJG and LRT wrote the original draft of the manuscript. All authors discussed the results and prepared and approved the final manuscript. LRT and CJG are the guarantors of this work and, as such, have full access to all the data in the study and take responsibility for the integrity of the data and the accuracy of the data analysis.

## Acknowledgments

We would like to acknowledge Dr. Ramiro Diz and the UNC Chapel Hill Flow Cytometry Core Facility; the UNC Chapel Hill Respiratory TRACTS Core Laboratory; Dr. Jake Dillard, Dr. Caitlyn Molloy, Dr. Nancie MacIver, and the MacIver Laboratory at UNC Chapel Hill; Nathan Spencer at syGlass, Inc.; and Benjamin Darwitz and the Thurlow Laboratory for advice and technical support.

## 8. Data Availability Statement

Data associated with three-dimensional analysis of germinal centers in vaccine-draining lymph nodes is made available through Zenodo in the dataset titled “Imaris files of germinal center surfaces from the lymph nodes of vaccinated insulin-deficient and control mice” (DOI: 10.5281/zenodo.17155425).

