## Supplemental Material for "Insulin-deficient diabetes impairs vaccine-mediated antibody and germinal center B-cell formation in mice"

**FOR**

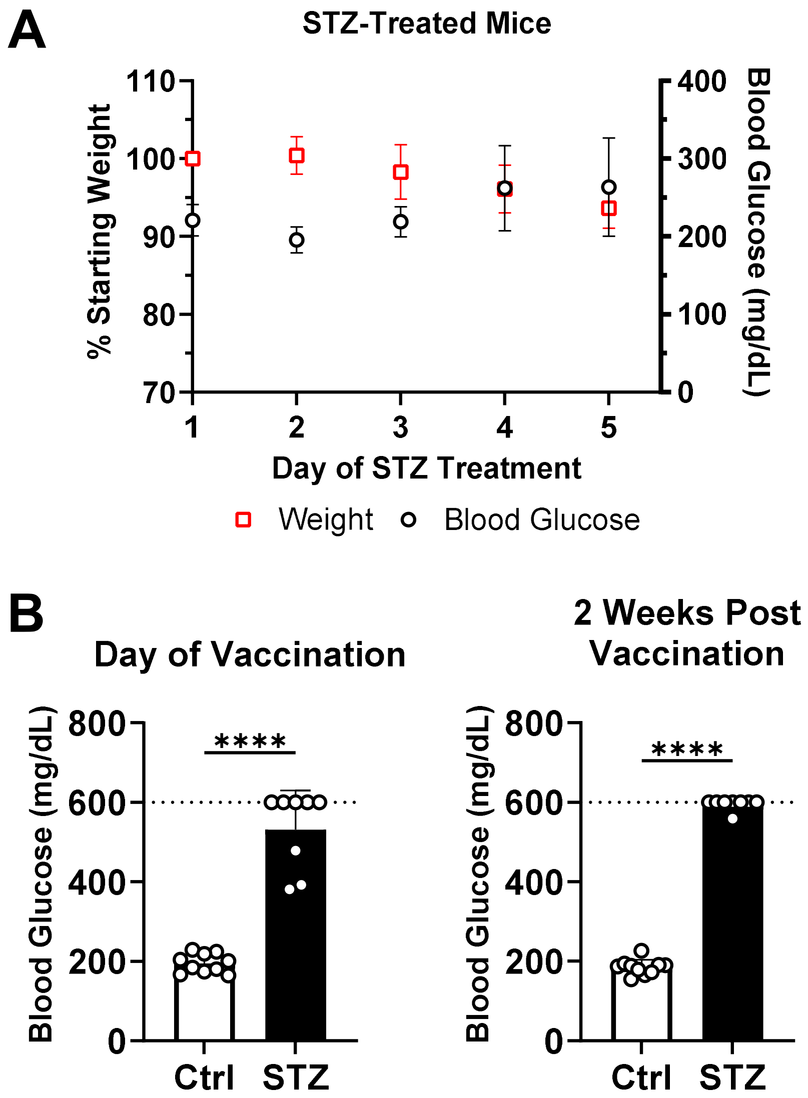


**Figure S1. Weight and elevated blood glucose during induction of diabetes. (A)** Mice were made insulin-deficient using streptozotocin (STZ) administered daily over a period of 5 days. Weight relative to the first day of STZ administration (red squares) and blood glucose (black circles) were measured each day of STZ administration (n = 10; data shown as mean ± SD). **(B)** Mice were made insulin-deficient using STZ. Immunizations were given 7 days after the last dose of STZ. Control mice (Ctrl), n = 10. STZ-treated mice, n = 8. Data is presented as mean + SD. Dotted line represents the limit of detection. Statistical comparisons were made using an unpaired *t*-test, ****$p < 0.0001$. Data is pooled from three independent experiments.

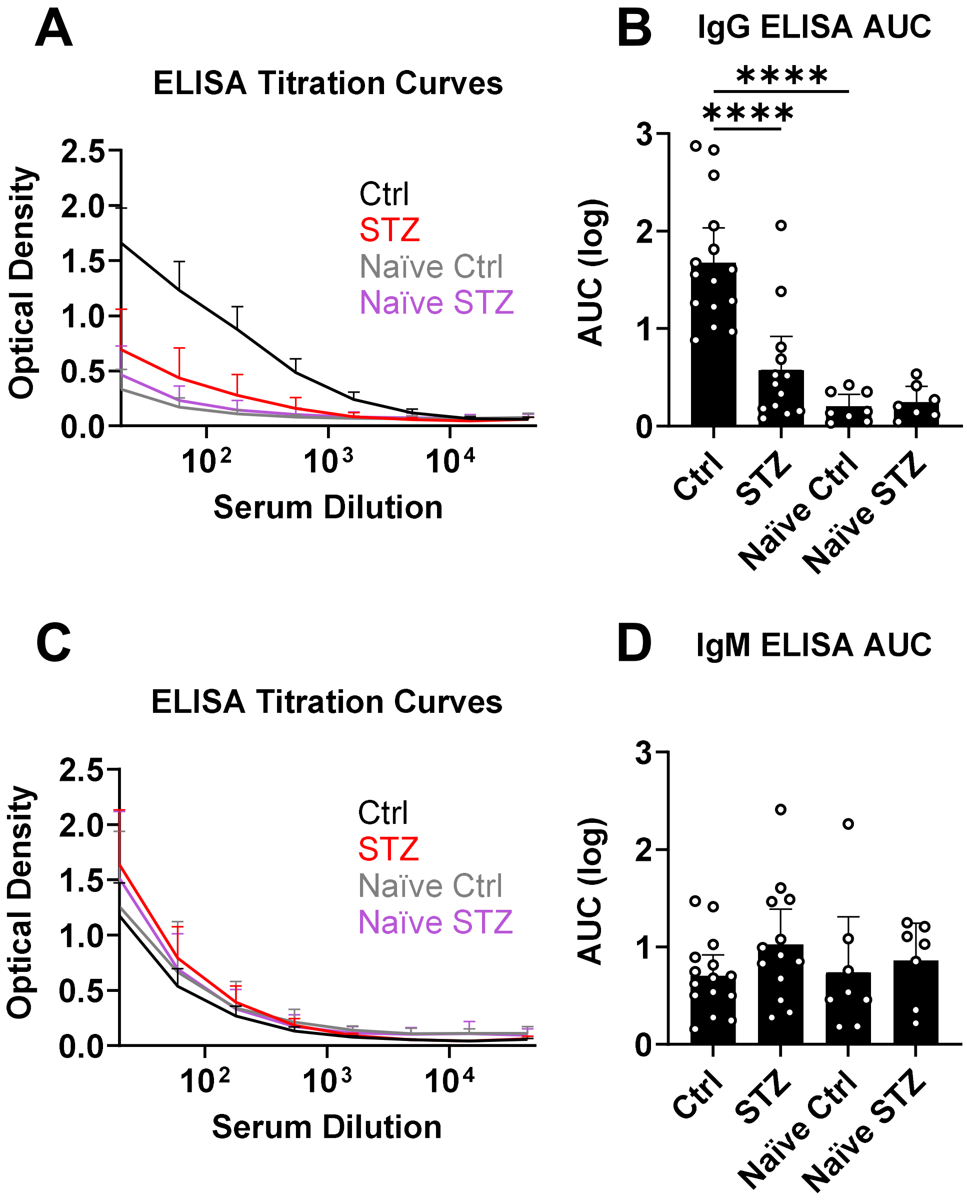


**Figure S2. Additional antibody measurements by ELISA.** Mice were made insulin-deficient using streptozotocin (STZ) and immunized with ovalbumin (Ova) adjuvanted with alum. Two weeks after immunization, Ova-specific **(A, B)** IgG and **(C, D)** IgM antibody levels were quantified by ELISA for control (Ctrl; n = 15), STZ-treated (n = 13), and naïve control (Naïve Ctrl, n = 8), and naïve STZ-treated (Naïve STZ, n = 7) mice. **(A, C)** Each group's mean (+95% clearance interval) optical density values (optical density at 450 nm wavelength corrected with 570 nm background subtraction) obtained from ELISA at each tested serum dilution. **(B, D)** Area under the curve (AUC) calculated for each animal's individual ELISA titration curve, using log-transformed serum dilutions. Data are presented as mean + 95% clearance interval. $^{****}p < 0.0001$, ANOVA with Tukey's test for multiple comparisons. Data is pooled from three independent experiments.

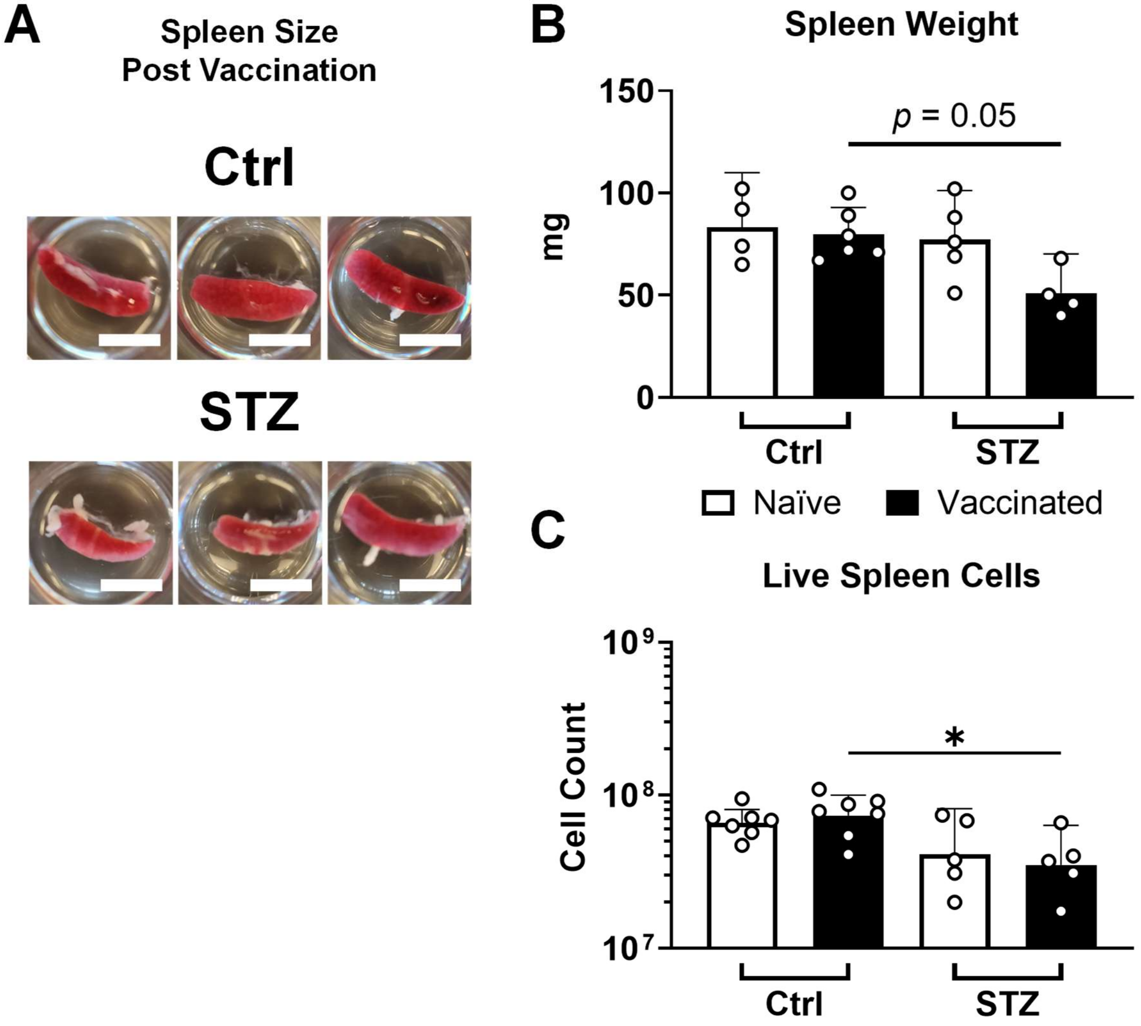


**Figure S3. Spleen size from vaccinated insulin-deficient mice.** Spleens were removed 2 weeks after vaccination with ovalbumin adjuvanted with alum from mice made insulin-deficient through streptozotocin administration (STZ) or from control animals (Ctrl). **(A)** Representative images of removed spleens. Scale bar = 15 mm. **(B)** Spleen weights (Ctrl: naïve n = 4, vaccinated n = 6; STZ: naïve n = 5, vaccinated n = 4). Data are represented as mean + 95% CI. **(C)** Cells were isolated from spleens and live cell counts were determined by staining with trypan blue (Ctrl: naïve n = 7, vaccinated n = 7; STZ: naïve n = 5, vaccinated n = 5). Data are represented as geometric mean + 95% CI. Statistical comparisons were made by ANOVA with Tukey's correction for multiple comparisons. $*p < 0.05$. Data was pooled from 2 independent experiments.

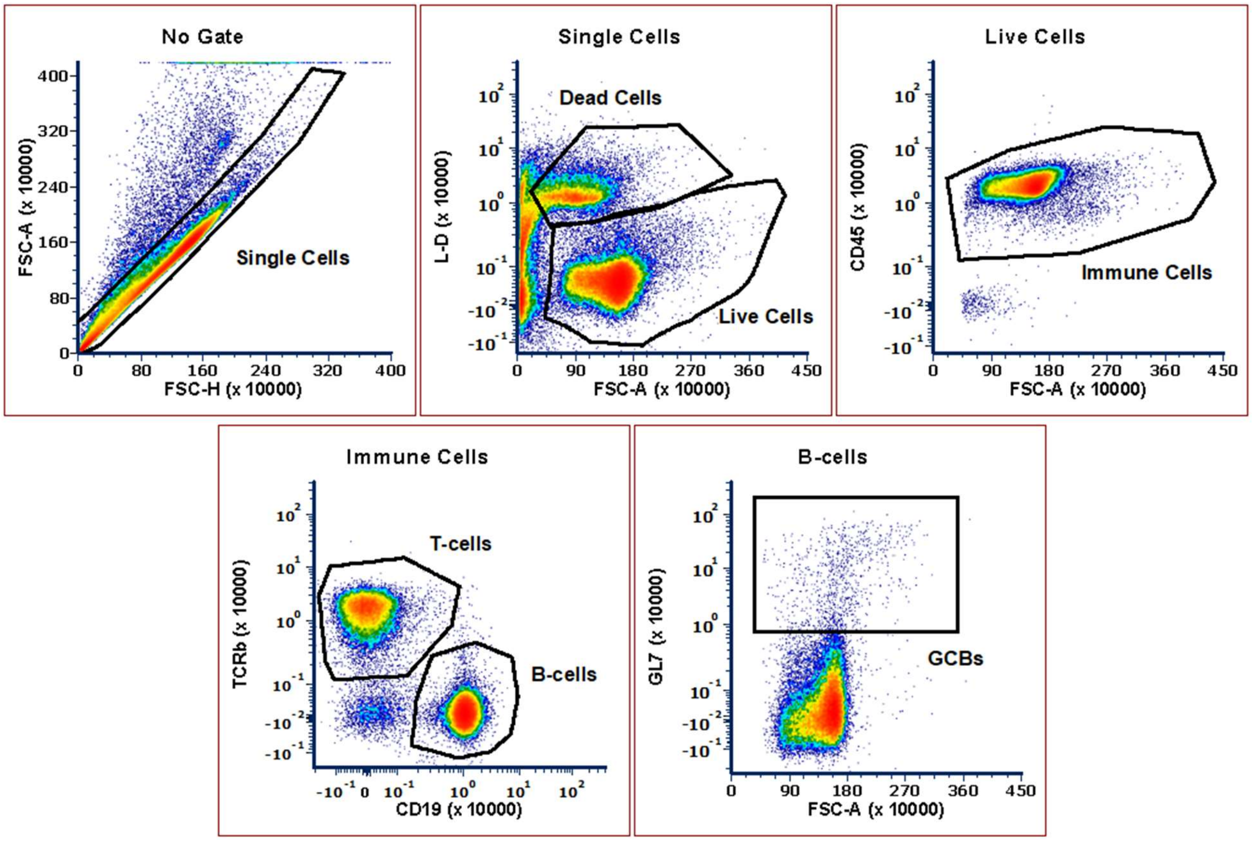


**Figure S4. Gating strategy for immune cell subset analysis by spectral flow cytometry.**

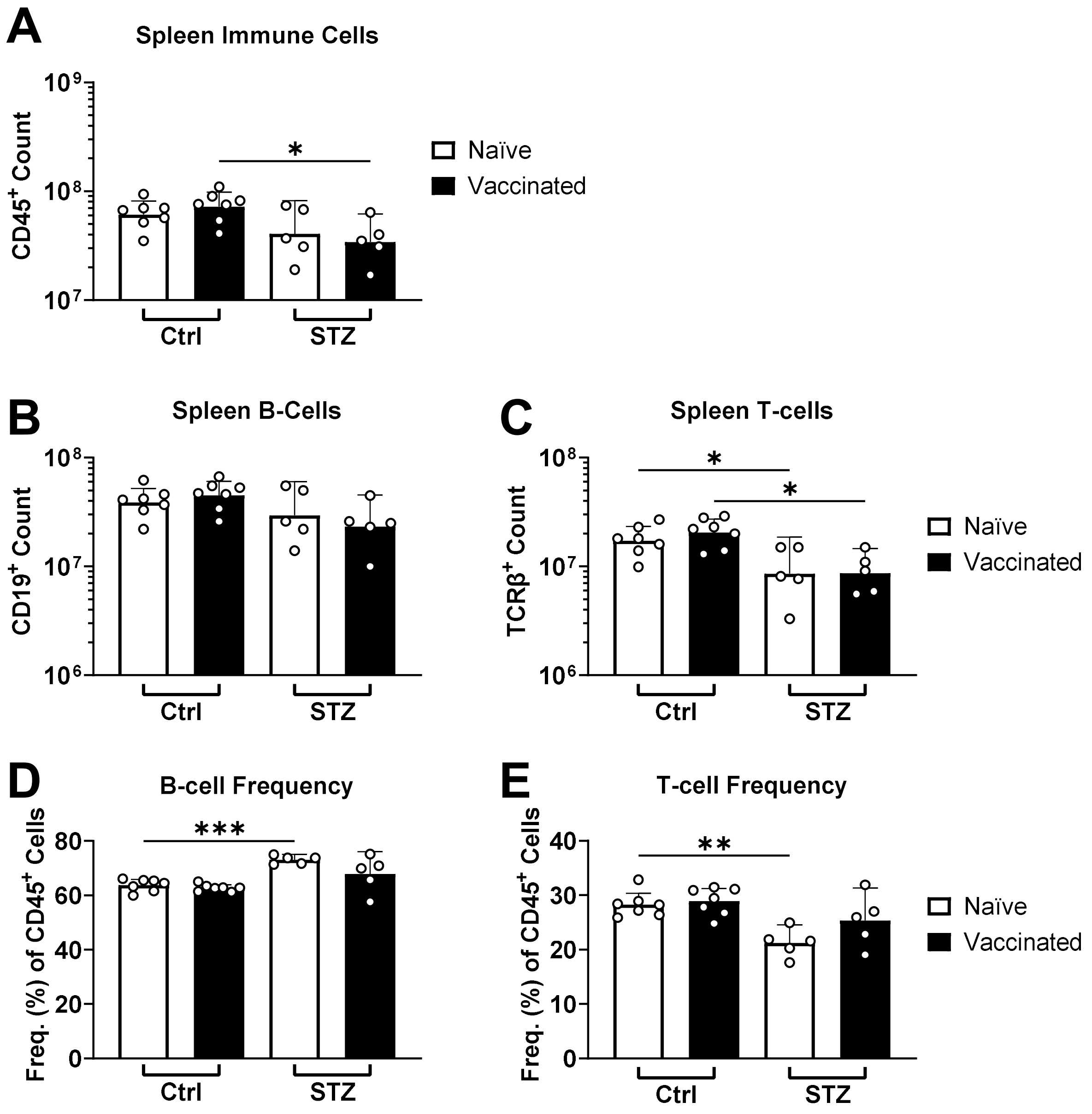


**Figure S5. Immune cell counts and frequencies in spleens after vaccination in insulin-deficient mice.** Total immune cell **(A)**, total B-cell **(B)**, and total T-cell **(C)** counts, as well as B-cell **(D)** and T-cell **(E)** frequencies among total immune cells, were quantified by flow cytometry in spleens 2 weeks after vaccination with ovalbumin adjuvanted with alum for insulin-deficient (STZ, naïve n = 5, vaccinated n = 5) and control (Ctrl, naïve n = 7, vaccinated n = 7) mice. Data is represented as geometric mean + 95% CI. Statistical comparisons were made using ANOVA with Tukey's correction for multiple comparisons. *$p < 0.05$, **$p < 0.01$, ***$p < 0.001$. Data was pooled from 2 independent experiments.

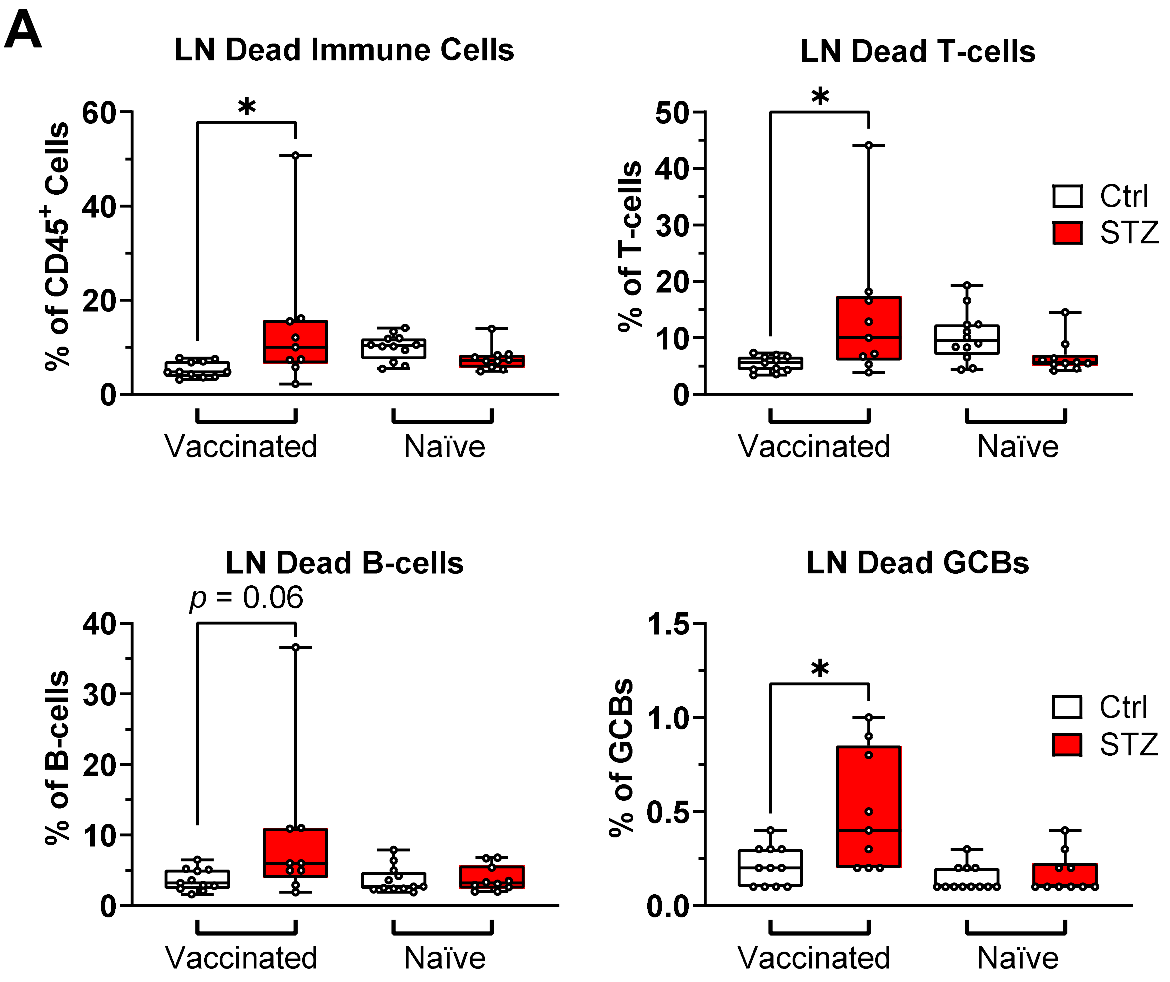
A
LN Dead Immune Cells
% of CD45+ Cells
*
Vaccinated
Naïve
LN Dead T-cells
% of T-cells
*
Ctrl
STZ
LN Dead B-cells
p = 0.06
% of B-cells
LN Dead GCBs
% of GCBs
*

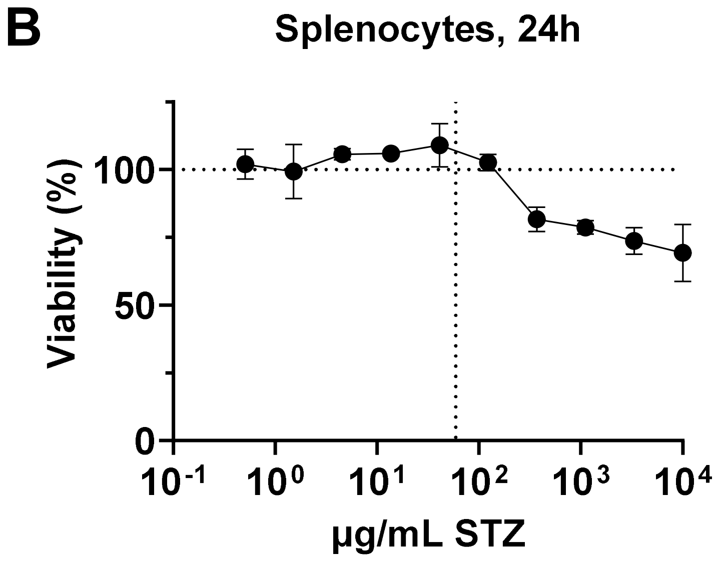
B
Splenocytes, 24h
Viability (%)
µg/mL STZ

**Figure S6. Streptozotocin treatment-related immune cell death. (A)** Frequency (%) of dead cells (viability-dye$^+$) in each immune cell population analyzed in draining lymph nodes (LN) 2 weeks after vaccination (ovalbumin and alum) by flow cytometry. Data are presented as quartiles with the center line at the median (control mice: naïve n = 12, vaccinated n = 11; STZ-treated mice: naïve n = 10, vaccinated n = 9). Vaccinated groups were statistically compared by Mann-Whitney *U* test. *$p < 0.05$. GCBs = germinal center B-cells. Data was pooled from 3 independent experiments. **(B)** Immune cells were isolated from the spleen of a control mouse and cultured *ex vivo* in the presence of streptozotocin (STZ) at various concentrations for 24h. Viability was determined by extracellular LDH detection, compared to untreated cells. Vertical dotted line represents the highest plasma concentration of STZ detected in mice 3 h after STZ administration reported by Anderson et al.[33] Each point represents mean and SD (n=3).

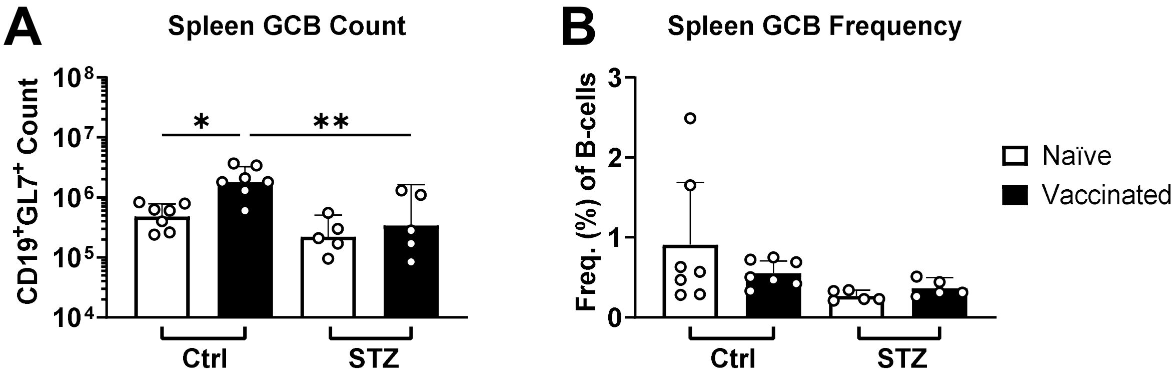


**Figure S7. Splenic germinal center B-cells (GCB) after vaccination in insulin-deficient mice.** GCB counts **(A)** and GCB frequencies **(B)** among total B-cells, were quantified by flow cytometry in spleens 2 weeks after vaccination with ovalbumin adjuvanted with alum for insulin-deficient (STZ, naïve n = 5, vaccinated n = 5) and control (Ctrl, naïve n = 7, vaccinated n = 7) mice. Data is represented as geometric mean + 95% CI. Statistical comparisons were made using ANOVA with Tukey's correction for multiple comparisons. *$p$ < 0.05, **$p$ < 0.01. Data was pooled from 3 independent experiments.

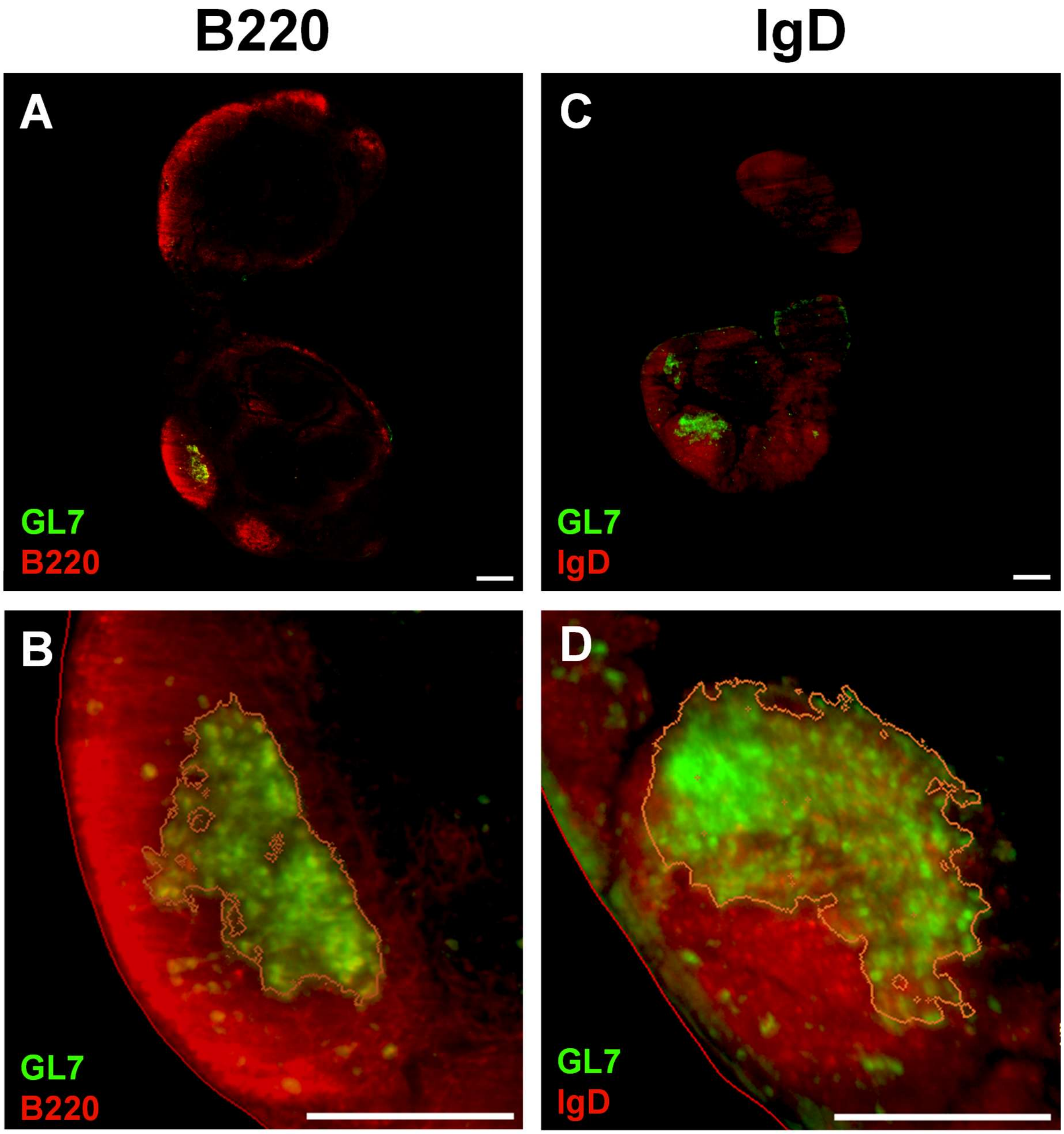


**Figure S8. B-cell follicle marker comparison.** 2D slice images from light sheet microscopy of draining inguinal lymph nodes 2 weeks after vaccination with ovalbumin adjuvanted with alum. Germinal center B-cells were detected by GL7 (green) and B-cell follicles (red) were detected either by B220 **(A, B)** or IgD **(C, D)**. In B and D, the red line represents the outside edge of the lymph node, and the orange line represents the edge of the germinal center as determined by analysis in syGlass. Scale bar 200 μm.

**Table S1. Antibodies and related conjugations used for spectral flow cytometry and light sheet microscopy.**

| Target | Conjugate | Vendor | Clone/ID | Use | Dilution |
|---|---|---|---|---|---|
| CD45 | PerCP/Cy5.5 | BioLegend | 30-F11 | Flow | 1:1600 |
| GL7 (Ly77) | PE | BioLegend | GL7 | Flow | 1:800 |
| CD19 | BV570 | BioLegend | 6D5 | Flow | 1:200 |
| TCRβ | BUV395 | BD | H57-597 | Flow | 1:100 |
| Rat IgG | AF647 | Jackson ImmunoResearch | 212-606-168 | Light sheet | 1:400 |
| B220 (CD45R) | Unlabeled | Invitrogen | RA3-6B2 | Light sheet | 1:800 |
| GL7 | Biotin | BioLegend | GL7 | Light sheet | 1:200 |
| Biotin (via streptavidin) | AF568 | Invitrogen | S11226A | Light sheet | 1:1000 |
| IgD | AF647 | BioLegend | 11-26c.2a | Light sheet | 1:100 |

**Table S2. Light sheet microscope settings per sample.** Chromatic correction was used. A-F = empty channel used to capture autofluorescence.

| Sample Name | Light Sheet Source | Number of Light Sheets | Laser Position Relative to Sample | Sheet NA | Sheet Width | Sheet Thickness (μm) | Zoom (X) | Z-spacing (μm) | Pixel Size (XY μm) | Cell Markers | | |
|---|---|---|---|---|---|---|---|---|---|---|---|---|
| | | | | | | | | | | 488 nm | 561 nm | 647 nm |
| **Ctrl 1** | Right | 3 | Middle | 0.039 | 100% | 7.1 | 3.2 | 3.55 | 0.95 | N/A | GL7 | B220 |
| **Ctrl 2** | Right | 3 | Middle | 0.032 | 100% | 7.1 | 2 | 5 | 1.54 | N/A | GL7 | B220 |
| **Ctrl 3** | Right | 3 | Middle | 0.024 | 100% | 11.5 | 2.5 | 5 | 1.22 | A-F | GL7 | B220 |
| **Ctrl 4** | Left | 3 | Middle | 0.024 | 100% | 11.5 | 2.5 | 5 | 1.22 | A-F | GL7 | IgD |
| **STZ 1** | Right | 3 | Slight right | 0.032 | 100% | 8.7 | 3.2 | 4.35 | 0.94 | A-F | GL7 | B220 |
| **STZ 2** | Right | 3 | Middle | 0.039 | 100% | 7.1 | 3.2 | 3.55 | 0.94 | A-F | GL7 | B220 |
| **STZ 3** | Right | 3 | Middle | 0.032 | 100% | 8.7 | 3.2 | 4.35 | 0.94 | A-F | GL7 | B220 |
| **STZ 4** | Left | 3 | Slight left | 0.032 | 100% | 8.7 | 3.2 | 4.35 | 0.94 | A-F | GL7 | B220 |

**Table S3. VR settings for GC identification.**

| Sample | Voxel Size (X-Y-Z µm) | Performance | Brightness | Contrast | Window min | Window max | Threshold min | Threshold max |
|---|---|---|---|---|---|---|---|---|
| **Ctrl 1** | 0.95. x 0.95 x 3.55 | 10.00 | 388.5 | 2.741 | 0 | 17863 | 1452 | 65535 |
| **Ctrl 2** | 1.54 x 1.54 x 5.00 | 10.00 | 8.256 | 1.035 | 0 | 40615 | 1254 | 65535 |
| **Ctrl 3** | 1.22 x 1.22 x 5.00 | 3.000 | 1.000 | 1.000 | 0 | 16191 | 827 | 65535 |
| **Ctrl 4** | 1.22 x 1.22 x 5.00 | 3.000 | 1.464 | 0.813 | 296 | 44204 | 699 | 65535 |
| **STZ 1** | 0.94 x 0.94 x 4.35 | 10.00 | 0.287 | 1.000 | 257 | 40394 | 2485 | 65535 |
| **STZ 2** | 0.94 x 0.94 x 3.55 | 7.753 | 38.62 | 1.438 | 2 | 28527 | 1601 | 65535 |
| **STZ 3** | 0.94 x 0.94 x 4.35 | 8.719 | 5.185 | 1.128 | 587 | 65535 | 1839 | 65535 |
| **STZ 4** | 0.94 x 0.94 x 4.35 | 3.000 | 12.55 | 1.231 | 0 | 65535 | 1741 | 65535 |
